# OpSeF: Open source Python framework for collaborative instance segmentation of bioimages

**DOI:** 10.1101/2020.04.29.068023

**Authors:** Tobias M. Rasse, Réka Hollandi, Péter Horváth

## Abstract

Various pre-trained deep learning models for the segmentation of bioimages have been made available as ‘developer-to-end-user’ solutions. They usually require neither knowledge of machine learning nor coding skills, are optimized for ease of use, and deployability on laptops. However, testing these tools individually is tedious and success is uncertain.

Here, we present the ‘Op’en ‘Se’gmentation ‘F’ramework (OpSeF), a Python framework for deep learning-based instance segmentation. OpSeF aims at facilitating the collaboration of biomedical users with experienced image analysts. It builds on the analysts’ knowledge in Python, machine learning, and workflow design to solve complex analysis tasks at any scale in a reproducible, well-documented way. OpSeF defines standard inputs and outputs, thereby facilitating modular workflow design and interoperability with other software. Users play an important role in problem definition, quality control, and manual refinement of results. All analyst tasks are optimized for deployment on Linux workstations or GPU clusters, all user tasks may be performed on any laptop in ImageJ.

OpSeF semi-automates preprocessing, convolutional neural network (CNN)-based segmentation in 2D or 3D, and post-processing. It facilitates benchmarking of multiple models in parallel. OpSeF streamlines the optimization of parameters for pre- and post-processing such, that an available model may frequently be used without retraining. Even if sufficiently good results are not achievable with this approach, intermediate results can inform the analysts in the selection of the most promising CNN-architecture in which the biomedical user might invest the effort of manually labeling training data.

We provide Jupyter notebooks that document sample workflows based on various image collections. Analysts may find these notebooks useful to illustrate common segmentation challenges, as they prepare the advanced user for gradually taking over some of their tasks and completing their projects independently. The notebooks may also be used to explore the analysis options available within OpSeF in an interactive way and to document and share final workflows.

Currently, three mechanistically distinct CNN-based segmentation methods, the U-Net implementation used in Cellprofiler 3.0, StarDist, and Cellpose have been integrated within OpSeF. The addition of new networks requires little, the addition of new models requires no coding skills. Thus, OpSeF might soon become both an interactive model repository, in which pre-trained models might be shared, evaluated, and reused with ease.

## Introduction

Phenomics, the assessment of the set of physical and biochemical properties that completely characterize an organism, has long been recognized as one of the greatest challenges in modern biology (Houle et al., 2010). Microscopy is a crucial technology to study phenotypic characteristics. Advances in high-throughput microscopy (Lang et al., 2006; Neumann et al., 2010; Chessel and Carazo Salas, 2019), slide scanner technology (Webster and Dunstan, 2014; Wang et al., 2019), light-sheet microscopy (Swoger et al., 2014; Ueda et al., 2020), semi-automated (Bykov et al., 2019; Schorb et al., 2019) and volume electron microscopy (Titze and Genoud, 2016; Vidavsky et al., 2016), as well as correlative light- and electron microscopy (Hoffman et al., 2020) have revolutionized the imaging of organisms, tissues, organoids, cells, and subcellular structures. Due to the massive amount of data produced by these approaches, the traditional biomedical image analysis tool of ‘visual inspection’ is no longer feasible, and classical, non-machine learning-based image analysis is often not robust enough to extract phenotypic characteristics reliably in a non-supervised manner.

Thus, the advances mentioned above were enabled by breakthroughs in the application of machine learning methods to biological images. Traditional machine learning techniques, based on random-forest classifiers and support vector machines, were made accessible to biologists with little to no knowledge in machine learning, using stand-alone tools such as ilastik (Haubold et al., 2016; Berg et al., 2019; Kreshuk and Zhang, 2019) or QuPath (Bankhead et al., 2017). Alternatively, they were integrated into several image analysis platforms such as Cellprofiler (Lamprecht et al., 2007), Cellprofiler Analyst (Jones et al., 2009), Icy (de Chaumont et al., 2012), ImageJ (Schneider et al., 2012; Arganda-Carreras et al., 2017) or KNIME (Sieb et al., 2007).

More recently, deep learning methods, initially developed for computer vision challenges, such as face recognition or autonomous cars, have been applied to biomedical image analysis (Cireşan et al., 2012; 2013). The U-Net is the most commonly used deep convolutional neural network specifically designed for semantic segmentation of biomedical images (Ronneberger et al., 2015). In the following years, neural networks were broadly applied to biomedical images (Zhang et al., 2015; Akram et al., 2016; Albarqouni et al., 2016; Çiçek et al., 2016; Milletari et al., 2016; Moeskops et al., 2016; Van Valen et al., 2016; Rajchl et al., 2017). Segmentation challenges like the 2018 Data Science Bowl (DSB) further promoted the adaptation of computer vision algorithms like Mask R-CNN (He et al., 2017) to biological analysis challenges (Caicedo et al., 2019). The DSB included various classes of nuclei. Schmidt and colleagues use the same dataset to demonstrate that star-convex polygons are better suited to represent densely packed cells (Schmidt et al., 2018b) than axis-aligned bounding boxes used in Mask R-CNN (Hollandi et al., 2020b). Training of deep learning models typically involves tedious annotation to create ground truth labels. Approaches that address this limitation include optimizing the annotation workflow by starting with reasonably good predictions (Hollandi et al., 2020a), applying specific preprocessing steps such that an existing model can be used (Whitehead, 2020), and the use of generalist algorithms trained on highly variable images (Stringer et al., 2020). Following the latter approach, Stringer and colleagues trained a neural network to predict vector flows generated by the reversible transformation of a highly diverse image collection. Their model includes a function to auto-estimate the scale. It works well on specialized and generalized data (Stringer et al., 2020).

Recently, various such pre-trained deep learning segmentation models have been published that are intended for non-machine learning experts in the field of biomedical image processing (Weigert et al., 2019; Hollandi et al., 2020b; Stringer et al., 2020). Testing such models on new data sets can be time-consuming and might not always give good results. Pre-trained models might fail because the new images do not resemble the data network was trained on sufficiently well. Alternatively, the underlying network architecture and specification, or the way data is internally represented and processed might not be suited for the presented task. Biomedical users with no background in computer science are often unable to distinguish these possibilities and might erroneously conclude that their problem is in principle not suited for deep learning-based segmentation. Thus, they might hesitate to create annotations to re-train the most appropriate architecture. Here, we present the **Op**en **Se**gmentation **F**ramework OpSeF, a Python framework for deep-learning-based instance segmentation of cells and nuclei. OpSeF has primarily been developed for staff image analysts with solid knowledge in image analysis, thorough understating of the principles of machine learning, and basic skills in Python. It wraps scikit-image, a collection of Python algorithms for image processing (van der Walt et al., 2014), the U-Net implementation used in Cellprofiler 3.0 (McQuin et al., 2018), StarDist (Schmidt et al., 2018b; Schmidt et al., 2018a; Weigert et al., 2019), and Cellpose (Stringer et al., 2020) in a single framework. OpSeF defines the standard in- and outputs, facilitates modular workflow design, and interoperability with other software. Moreover, it streamlines and semi-automates preprocessing, CNN-based segmentation, postprocessing as well as evaluation of results. Jupyter notebooks (Kluyver et al., 2016) serve as a minimal graphical user interface. Most computations are performed head-less and can be executed on local workstations as well as on GPU clusters. Segmentation results can be easily imported and refined in ImageJ using AnnotatorJ (Hollandi et al., 2020a).

## Material and Methods

### Data description

#### Cobblestones

Images of cobblestones were taken with a Samsung Galaxy S6 Active Smartphone.

#### Leaves

Noise was added to the demo data from ‘YAPiC - Yet Another Pixel Classifier’ available at https://github.com/yapic/yapic/tree/master/docs/example_data using the *Add Noise* function in ImageJ.

#### Small fluorescent nuclei

Images of Hek293 human embryonic kidney stained with a nuclear dye from the image set BBBC038v1(Caicedo et al., 2019) available from the Broad Bioimage Benchmark Collection (BBBC) were used. Metadata is not available for this image set to confirm staining conditions. Images were rescaled from 360×360 pixels to 512×512 pixels.

#### 3D colon tissue

We used the low signal-to-noise variant of the image set BBBC027 (Svoboda et al., 2011) from the BBBC showing 3D colon tissue images.

#### Epithelial cells

Images of cervical cells from the image set BBBC038v1 (Caicedo et al., 2019) available from the BBBC display cells stained with a dye that labels membranes weakly and nuclei strongly. The staining pattern is reminiscent of images of methylene blue-stained cells. However, metadata is not available for this image set to confirm staining conditions.

#### Skeletal muscle

A methylene blue-stained skeletal muscle section was recorded on a Nikon Eclipse Ni-E microscope equipped with a Märzhäuser SlideExpress2 system for automated handling of slides. The pixel size is 0.37*0.37 μm. Thirteen large patches of 2048×2048 pixels size were manually extracted from the original 44712×55444 pixels large image. Color images were converted to greyscale.

#### Kidney

HE stained kidney paraffin sections were recorded on a Nikon Eclipse Ni-E microscope equipped with a Märzhäuser SlideExpress2 system for automated handling of slides. The pixel size is 180 × 180 nm. The original, stitched, 34816 × 51200 pixels large image was split into two large patches (18432 × 6144 and 22528 × 5120 pixel). Next, the Eosin staining was extracted using the *Color Deconvolution* ImageJ plugin. This plugin implements the method described by Ruifrok and Johnston (Ruifrok and Johnston, 2001).

#### Arabidopsis flowers

H2B:mRuby2 was used for the visualization of somatic nuclei of *Arabidopsis thaliana* flower. The flower was scanned from eight views differing by 45° increments in a Zeiss Z1 light-sheet microscope (Valuchova et al., 2020). We used a single view to mimic a challenging 3D segmentation problem. Image files are available in the Image Data Resource (Williams et al., 2017) under the accession code: idr0077.

#### Mouse blastocysts

The DAPI signal from densely packed E3.5 mouse blastocysts nuclei was recorded on a Leica SP8 confocal microscope using a 40x 1.30 NA oil objective (Blin et al., 2019). Image files are available in the Image Data Resource (Williams et al., 2017) under the accession code: idr0062.

#### Neural monolayer

The DAPI signal of a neural monolayer was recorded on a Leica SpE confocal microscope using a 63x 1.30 NA oil objective (Blin et al., 2019). Image files are available in the Image Data Resource (Williams et al., 2017) under the accession code: idr0062.

### Algorithm

Ideally, OpSeF is used as part of collaborative image analysis projects, to which both the user and the image analyst contribute their unique expertise (**Figure 1A**). If challenges arise, the image analyst **(Figure 1A)** might consult other OpSeF users or the developer of tools used within OpSeF. The analyst will – to the benefit of future users – become more skilled using CNN-based segmentation in analysis workflows. The user, who knows the sample best, plays an important role in validating results and discovering artifacts **(Figure 1A)**. Exemplary workflows and new models might be shared to the benefit of other OpSeF users (**Figure 1B**).

**Figure 1.**
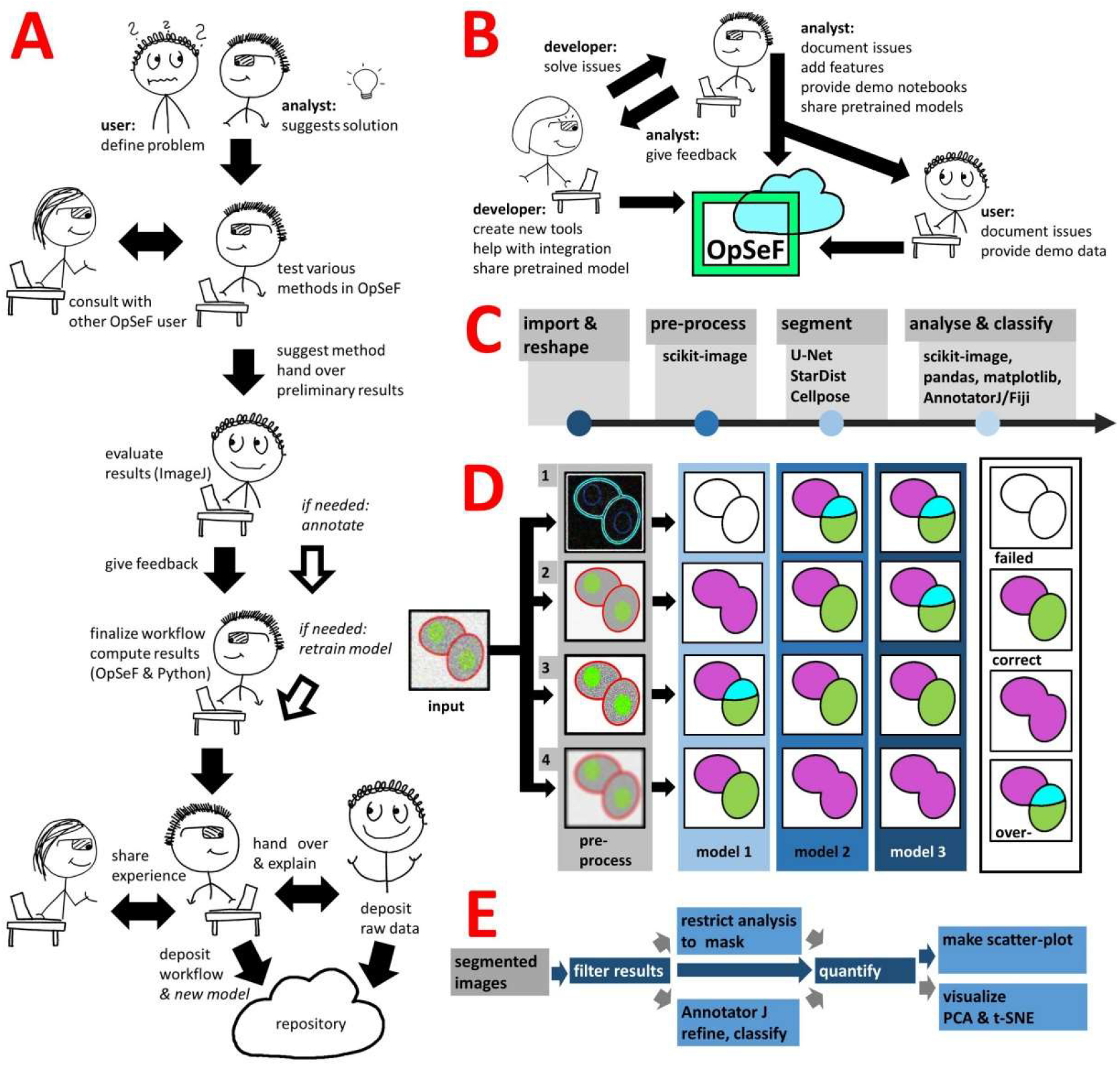
Image analysis using OpSeF **(A)** Illustration on how to use OpSeF for collaborative image analysis **(B)** Illustration on how developers, image analysts and users might contribute towards the further development of OpSeF. **(C)** OpSeF analysis pipeline consists of four groups of functions used to *import and reshape* the data, to *preprocess* it, to *segment objects*, and to *analyze and classify* results. **(D)** Optimization procedure. Left panel: Illustration of a processing pipeline, in which three different models are applied to data generated by four different preprocessing pipelines each. Right panel: Resulting images are classified into results that are correct; suffer from under- or over-segmentation or fail to detect objects. **(E)** Illustration of post-processing pipeline. Segmented objects might be filtered by their region properties or a mask, results might be exported to AnnotatorJ and re-imported for further analysis. Blue arrows define the default processing pipeline, grey arrows feature available options. Dark blue boxes are core components, light blue boxes are optional processing steps.

OpSeF’s analysis pipeline consists of four principal sets of functions to *import and reshape* the data, to *preprocess* it, to *segment* objects, and to *analyze and classify* results (**Figure 1C**). Currently, OpSeF can process individual tiff files and the proprietary Leica ‘.lif’ container file format. During *import and reshape*, the following options are available for tiff-input: *tile* in 2D and 3D, *scale*, and make *sub-stacks*. For lif-files, only the make *sub-stacks* option is supported. Preprocessing is mainly based on scikit-image (van der Walt et al., 2014). It consists of a linear workflow in which 2D images are filtered, the background is removed, and stacks are projected. Next, the following optional preprocessing operations might be performed: histogram adjustment (Zuiderveld, 1994), edge enhancement, and inversion of images. Available segmentation options include the pretrained U-Net used in Cellprofiler 3.0 (McQuin et al., 2018), the StarDist 2D model (Schmidt et al., 2018b; Schmidt et al., 2018a) and Cellpose (Stringer et al., 2020). Available options for preprocessing in 3D are limited (Figure 1B, lower panel). Segmentation in 3D is computationally more demanding. Thus, we recommend a two-stage strategy. Preprocessing parameters are first explored on representative planes in 2D. Next, further optimization in 3D is performed. Either way, preprocessing and selection of the ideal model for segmentation are one functional unit. **Figure 1D** illustrates this concept with a processing pipeline, in which three different models are applied to four different preprocessing pipelines each. The resulting images are classified into results that are mostly correct, suffer from under- or over-segmentation, or largely fail to detect objects. In the given example, the combination of preprocessing pipeline three and model two gives overall the best result. We recommend an iterative optimization which starts with a large number of models, and relatively few, but conceptually distinct preprocessing pipelines. For most datasets some models outperform others. In this case, we recommend fine-tuning the most promising preprocessing pipelines in combination with the most promising model. OpSeF uses matplotlib (Virtanen et al., 2020) to visualize results in Jupyter notebooks and to export exemplary results that may be used as figures in publications. All data is managed in pandas (Virtanen et al., 2020) and might be exported as csv file. Scikit-image (van der Walt et al., 2014), and scikit-learn (Pedregosa et al., 2011) (**Figure 1E** and **Figure 2A-C**) are used for pre- and postprocessing of segmentation results, which might e.g. be filtered based on their size, shape or other object properties (**Figure 2B**). Segmentation objects may further be refined by a user-provided (**Figure 2D**) or an autogenerated mask. Results might be exported to AnnotatorJ (Hollandi et al., 2020a) for editing or classification in ImageJ. AnnotatorJ is an ImageJ plugin that helps hand-labeling data with deep learning-supported semi-automatic annotation and further convenient functions to easily create and edit object contours. It has been extended with a classification mode and import/export fitting the data structure used in OpSeF. After refinement, results can be re-imported and further analyzed in OpSeF. Analysis options include scatter plots of region properties (**Figure 2B**), T-distributed Stochastic Neighbor Embedding (t-SNE) analysis (**Figure 2F**) and principal component analysis (PCA) (**Figure 2G**).

**Figure 2.**
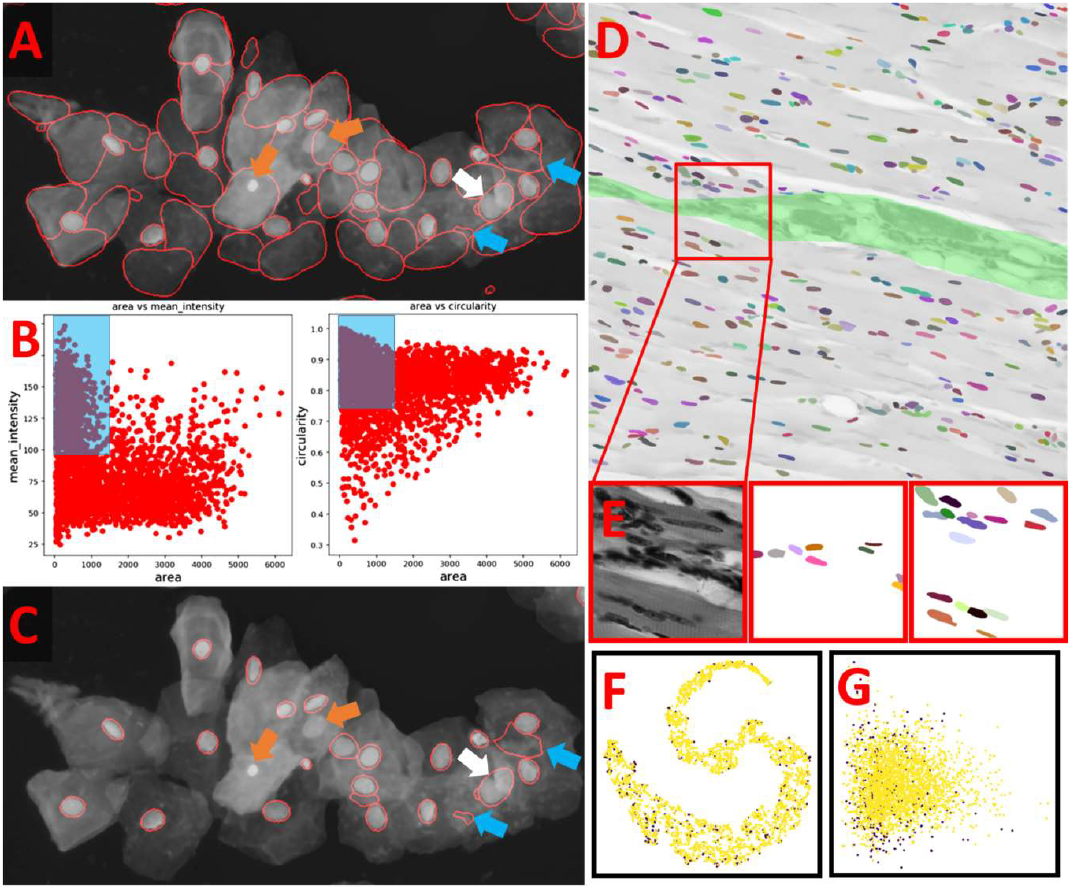
Example of how post-processing can be used to refine results **(A)** StarDist segmentation of the multi-labeled cells dataset detected nuclei reliably but caused many false positive detections. These resemble the typical shape of cells but are larger than true nuclei. Orange arrows point at nuclei that were missed, the white arrow at two nuclei that were not split, the blue arrows at false positive detections that could not be removed by filtering. **(B)** Scatter plot of segmentation results shown in **A**. Left panel: Mean intensity plotted against object area. Right panel: Circularity plotted against object area. Blue Box illustrates the parameter used to filter results. **(C)** Filtered Results. Orange arrows point at nuclei that were missed, the white arrow at two nuclei that were not split, the blue arrows at false positive detections that could not be removed by filtering. **(D,E)** Example for the use of a user-provided mask to classify segmented objects. The segmentation results (false-colored nuclei) are superimposed onto the original image subjected to [median 3×3] preprocessing. All nuclei located in the green area are assigned to Class 1, all others to Class 2. The red box indicates the region shown enlarged in **(E)**. From left to right in **E**: original image, nuclei assigned to class 1, nuclei assigned to class 2. **(F)** T-distributed Stochastic Neighbor Embedding (t-SNE) analysis of nuclei assigned to class 1 (purple) or class 2 (yellow). **(G)** Principal component analysis (PCA) of nuclei assigned to class 1 (purple) or class 2 (yellow).

### Data and Source code availability

Test data for the Jupyter notebooks is available at: https://owncloud.gwdg.de/index.php/s/nSUqVXkkfUDPG5b

Test data for AnnotatorJ is available at: https://owncloud.gwdg.de/index.php/s/dUMM6JRXsuhTncS

The source code is available at: https://github.com/trasse/OpSeF-IV

## Results

We provide demonstration notebooks to illustrate how OpSeF might be used to elucidate efficiently whether a given segmentation task is solvable with state of the art deep convolutional neural networks (CNNs). In the first step preprocessing parameters are optimized. Next, we test whether the chosen model performs well without re-training. Finally, we assess how well it generalizes on heterogeneous datasets.

Preprocessing can be used to make the input image more closely resemble the visual appearance of the data on which the models for the CNNs bundled with OpSeF were trained on e.g. by filtering and resizing. Additionally, preprocessing steps can be used to normalize data and reduce heterogeneity. Generally, there is not a single, universally best preprocessing pipeline. Instead, well-performing combinations of preprocessing pipelines and matching CNN-models can be identified. Even the definition of a ‘good’ result depends on the biological question posed and may vary from project to project. For cell tracking, very reproducible cell identification will be of utmost importance; for other applications, the accuracy of the outline might be more crucial. To harness the full power of CNN-based segmentation models and to build trust in their more widespread use, it is essential to understand under which conditions they are prone to fail.

We use various demo datasets to challenge the CNN-based segmentation pipelines. Jupyter notebooks document how OpSeF was used to obtain reliable results. These notebooks are provided as a starting point for the iterative optimization of user projects and as a tool for interactive user training.

The first two datasets – cobblestones and leaves – are generic, non-microscopic image collections, designed to illustrate common analysis challenges. Further datasets exemplify the segmentation of a monolayer of fluorescent cells, fluorescent tissues, cells in which various compartments have been stained with the same dye, as well as histological sections stained with one or two dyes. The latter dataset exemplifies additionally how OpSeF can be used to process large 2D images.

Nuclei and cells used to train CNN-based segmentation are most commonly round or ellipsoid shaped. Objects in the cobblestone dataset are approximately square-shaped. Thus, the notebook may be used as an example to train segmentation of non-round cells (e.g. many plant cells, neurons). Heterogeneous intensities within objects and in the border region, as well as a five-fold variation of object size, further challenge segmentation pipelines. In the first round of optimization, minor smoothing [median filter with 3×3 kernel (median 3×3)] and background subtraction were applied. Next, the effect of additional histogram equalization, edge enhancement, and image inversion was tested. The resulting four preprocessed images were segmented with all models [Cellpose nuclei, Cellpose Cyto, StarDist, and U-Net]. The Cellpose scale-factor range [0.2,0.4,0.6] was explored. Among the 32 resulting segmentation pipelines, the combination of image inversion and the Cellpose Cyto 0.4 model produced the best results in both training images (**Figure 3A,B**) without further optimization. The segmentation generalized well to the entire dataset. Only in one image, three objects were missed, and one object was over-segmented. Borders around these stones are very hard to distinguish for a human observer, and even further training might not resolve the presented segmentation tasks (**Figure 3E,F**). Cellpose has been trained on a large variety of images and had been reported to perform well on objects of similar shape (compare Figure 4, Images 21,22,27 in (Stringer et al., 2020)).

**Figure 3.**
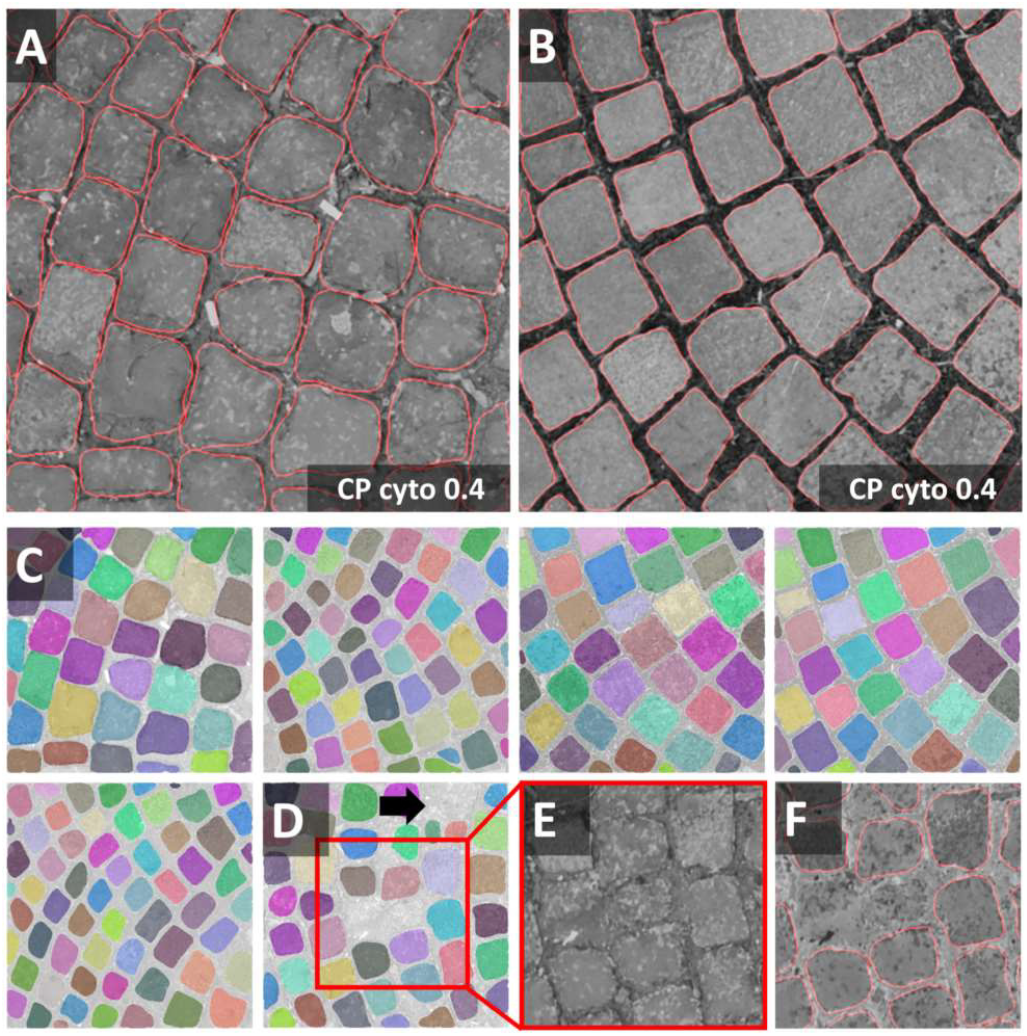
Cobblestones notebook: Segmentation of non-roundish cells **(A,B)** Segmentations (red line) of the Cellpose Cyto 0.4 model are superimposed onto the original image subjected to [median 3×3] preprocessing. The inverted image (not shown) was used as input to the segmentation. Outlines are well defined, no objects were missed, none over-segmented. These settings fit accurately to the entire dataset (train and test) shown in **(C,D)**. Only in one image, three objects were missed and one was over-segmented. Borders around these stones are hard to discern. Individual objects are false color-coded in **C,D**. The red squares in **D** highlight one of the two problematic regions shown as a close-up in **(E)** and (**F)**.

**Figure 4.**
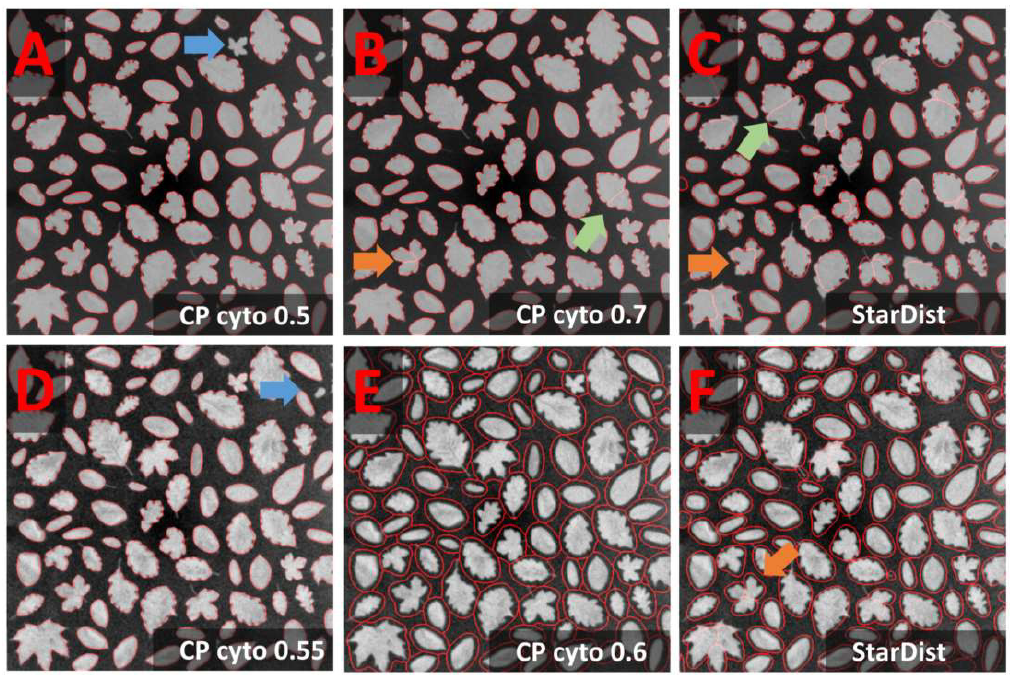
Leaves notebook: Segmentation of rare concave cells **(A-C)** Segmentation results (red line) of the Cellpose Cyto 0.5 and 0.7 model are superimposed onto the original image subjected to [median 3×3] preprocessing. The inverted image (not shown) was used as input to the segmentation. Outlines are well defined, few objects were missed (**A:** blue arrow), some over-segmented (**B,C:** green and orange arrow). Green arrow points at an oak leave with prominent leaf veins, orange arrow at maple leaves with less prominent leave veins. **(D-F)** Results of further optimization. Further smoothing **(E)** reduced the rate of false negatives (blue arrow) and over-segmentation in the Cellpose Cyto model. However, object outlines were less precise **(E)**.

Segmentation of the leaves dataset seems trivial and could easily be solved by any threshold-based approach. Nevertheless, it challenges CNN-based segmentation due to the presence of concave and covex shapes. Thus, it is ideal to illustrate the segmentation of amoeboid cells. Moreover, objects contain dark lines, vary 20-fold in area, and are presented on a heterogeneous background. Preprocessing was performed as described for the cobblestone dataset. The most promising result was obtained with the Cellpose Cyto 0.5 model in combination with [median 3×3 & image inversion] preprocessing (**Figure 4A,B**) and the StarDist model with [median 3×3 & histogram equalization] preprocessing (**Figure 4C**). Outlines were well defined, few objects were missed (blue arrow in **Figure 4A**), few over-segmented (green and orange arrow in **Figure 4B,C**). The Cellpose Cyto 0.7 model gave similar results.

Maple leaves (orange arrows in **Figure 4B,C**) were most frequently over-segmented. Their shape resembles a cluster of touching cells. Thus, the observed over-segmentation might be caused by the attempt of the CNN to reconcile their shapes with structures it has been trained on. Oak leaves were the second most frequently over-segmented leaf type. These leaves contain dark leaf veins that might be interpreted as cell borders. However, erroneous segmentation mostly does not follow these veins (green arrow in **Figure 4B**). Next, the effect of stronger smoothing [mean 7×7] was explored. The Cellpose nuclei model (**Figure 4E**) reduced the rate of false-negative detections (**Figure 4D** blue arrow) and over-segmentation (**Figure 4F** orange arrow) at the expense of loss in precision of object outlines. Parameter combinations tested in **Figure 4D,E** generalize well in the entire dataset.

Next, we used OpSeF to segment nuclei in a monolayer of cells. Most nuclei are well separated. We focused our analysis on the few touching nuclei. Both the Cellpose nuclei model and the Cellpose Cyto model performed well across a broad range of scale-factors. Interestingly, strong smoothing made the Cellpose nuclei but not the Cellpose Cyto model more prone to over-segmentation (**Figure 5A**). The StarDist model performed well, while the U-Net failed surprisingly, given the seemingly simple task. Pixel intensities have a small dynamic range, and nuclei are dim and rather large. To elucidate whether any of these issues led to this poor performance, we binned the input 2×2 (U-Net+BIN panel in **Figure 5A**) and adjusted brightness and contrast. Adjusting brightness and contrast alone had no beneficial effect (data not shown). The U-Net performed much better on the binned input. Subsequently, we batch-processed the entire dataset. StarDist was more prone to over-segmentation (green arrow in **Figure 5B**), but detected smaller objects more faithfully (orange arrow in **Figure 5B**) and was more likely to include atypical objects, e.g. nuclei during cell division that display a strong texture (blue arrow in **Figure 5B**). Substantial variation in brightness was well tolerated by both models (white arrow in **Figure 5B**). Both models complement each other well.

**Figure 5.**
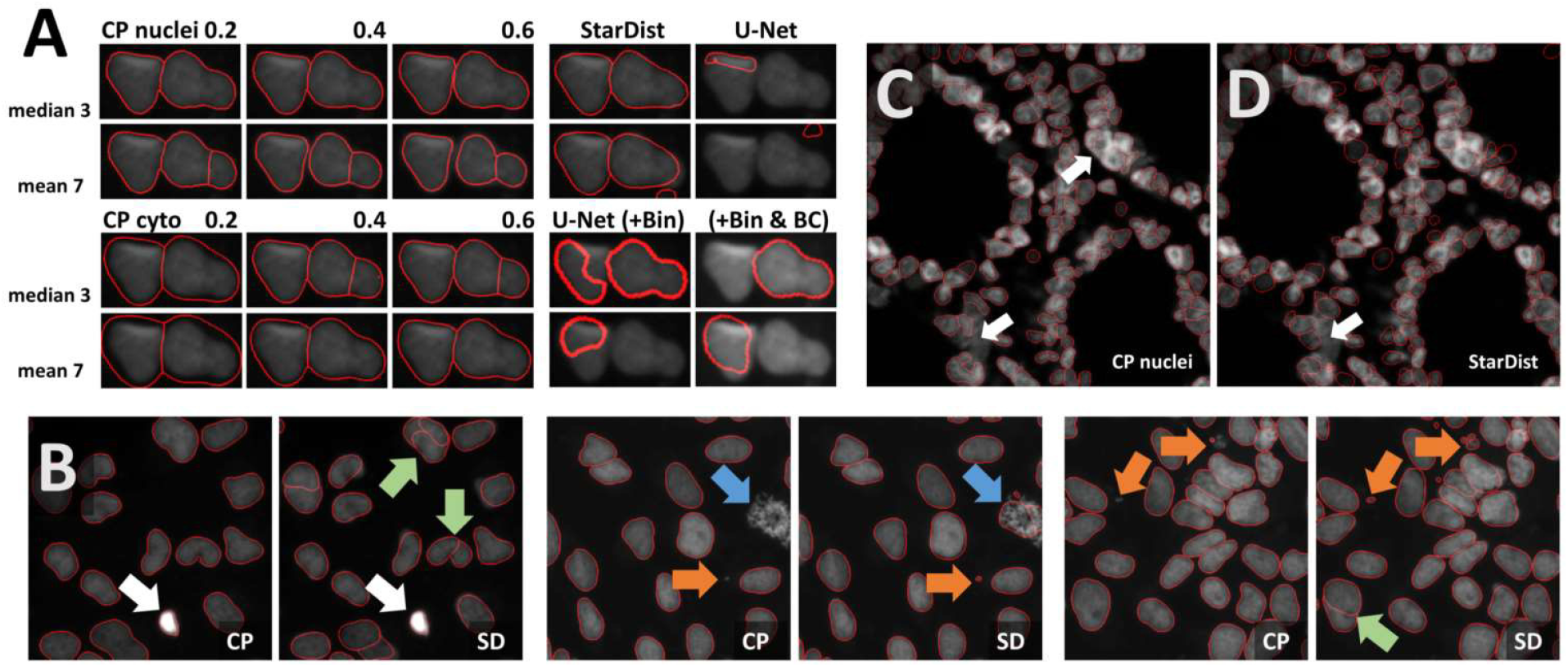
Segmentation of dense fluorescent nuclei **(A)** Segmentation of the sparse nuclei data set. Example of two touching cells. The segmentation result (red line) superimposed onto the original image subjected to [median 3×3] preprocessing. Columns define the model used for segmentation, the rows the filter used for preprocessing. Cellpose nuclei model and the Cellpose Cyto model performed well across a large range of scale-factors. The U-Net failed initially. Improved results after binning are shown in the lower right panel. **(B)** Comparison of Cellpose nuclei (CP) and StarDist (SD) segmentation results (red line). StarDist is more prone to over-segmentation (green arrow), but detects smaller objects more reliably (orange arrow) and tends to include objects with strong texture (blue arrow). Strong variation in brightness was tolerated well by both models (white arrow). **(C,D)** Segmentation of 3D colon tissue from the Broad BioImage Benchmark Collection with Cellpose nuclei (CP) or StarDist (SD). Both models gave reasonable results. Only a few dense clusters could not be segmented (white arrow).

We also tested a more complex dataset: 3D colon tissue from the Broad Bioimage Benchmark Collection. This synthetic dataset is ideally suited to assess segmenting clustered nuclei in tissues. We chose the low signal-to-noise variant, which allowed us to test denoising strategies. Sum, maximum, and median projection of three Z-planes was tested in combination with the preprocessing variants previously described for the monolayer of cells dataset. Twelve different preprocessing pipelines were combined with all models [Cellpose nuclei, Cellpose Cyto, StarDist, and U-Net]. The Cellpose scale-factor range [0.15, 0.25, 0.4, 0.6] was explored. Many segmentation decisions in the 3D colon tissue dataset are hard to perform even for human experts. Within this limitation, [median projection & histogram equalization] preprocessing produced reasonable results without any further optimization in combination with either Cellpose nuclei 0.4 or the StarDist model (**Figure 5C,D**). Only a few cell clusters were not segmented (**Figure 5C,D** white arrow). Both models performed equally well on the entire data set.

We subsequently tried to segment a single layer of irregular-shaped epithelial cells, in which the nucleus and cell membranes had been stained with the same dye. In the first run, minor [median 3×3] or strong [mean 7×7] smoothing was applied. Next, the effect of additional histogram equalization, edge enhancement, and image inversion was tested. The resulting eight preprocessed images were segmented with all models [Cellpose nuclei, Cellpose Cyto, StarDist, and U-Net]. The Cellpose scale-factor range [0.6, 0.8, 1.0, 1.4, 1.8] was explored. The size of nuclei varied more than five-fold. We thus focused our analysis on a cluster of particularly large nuclei and a cluster of small nuclei. The Cellpose nuclei 1.4 and StarDist model detected both small and large nuclei similarly well (**Figure 6A**). StarDist segmentation results included many cell-shaped false positive detections. As they were much larger than true nuclei, they could be filtered out during post-processing. While the U-Net did not perform well on the same input [median 3×3] (**Figure 6A**), it returned better results (**Figure 6A**) upon [mean 7×7 & histogram equalization] preprocessing. As weak smoothing was beneficial for the Cellpose and StarDist pipelines and stronger smoothing for the U-Net pipelines, we explored the effect of intermediate smoothing [median 5×5] for Cellpose and StarDist and even stronger smoothing [mean 9×9] for the U-Net pipelines. A slight improvement was observed. Thus, we used [median 5×5] preprocessing in combination with Cellpose nuclei 1.5 or StarDist model to process the entire dataset. Cellpose frequently failed to detect bright, round nuclei (**Figure 6B**, arrows) and StartDist had many false detections. Thus, re-training or post-processing is required.

**Figure 6.**
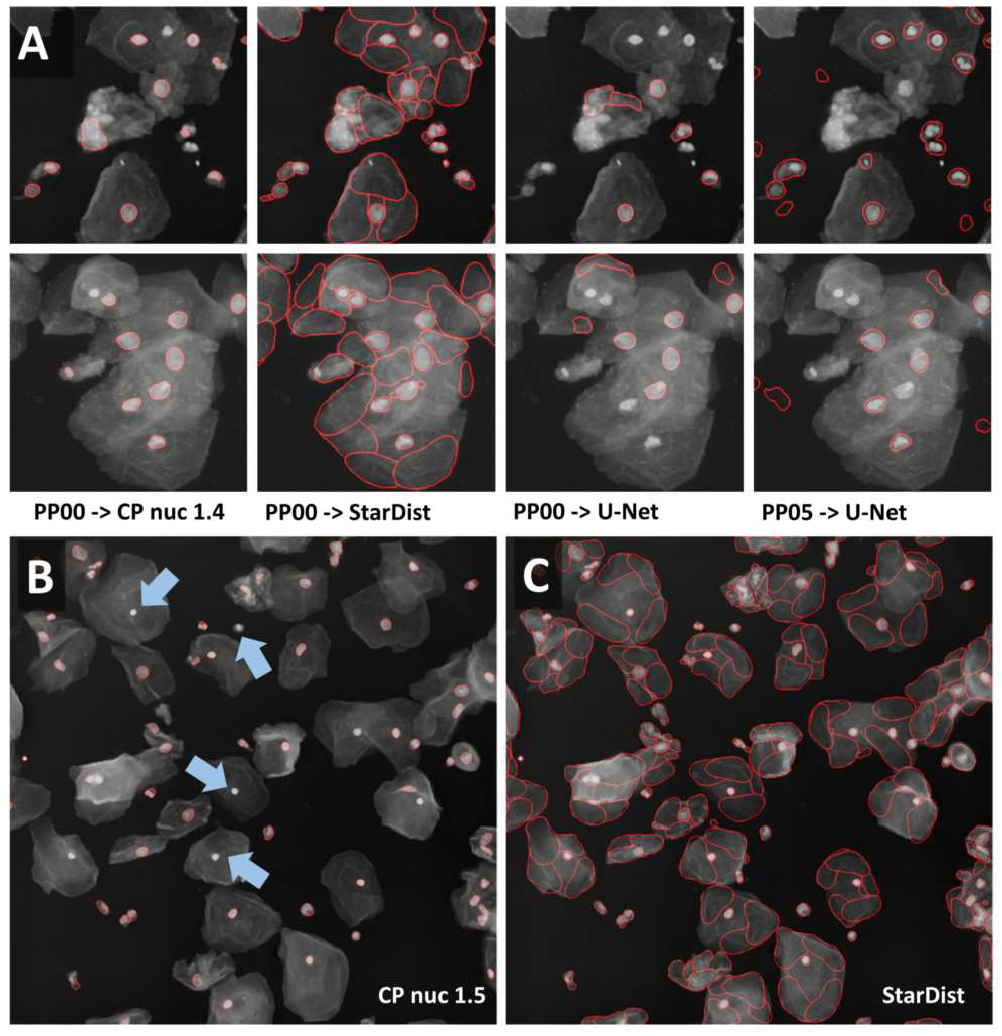
Segmentation of multi-labeled cells **(A)** Images of large epithelial cells from the 2018 Data Science Bowl collection were used to test the segmentation of a single layer of cells. These cells vary in size. The Cellpose nuclei 1.4 model and [median 3×3] preprocessing gave a reasonable segmentation for large and small nuclei. StarDist segmentation based on the same input detected nuclei more reliably. However, many false-positive detections were present. Interestingly, the shape of false detections resembles the typical shape of cells well. The U-Net did not perform well with the same preprocessing, but with [mean 7×7 & histogram equalization] preprocessing. **(B)** [Median 5×5] preprocessing in combination with the Cellpose 1.5 nuclei or the StarDist model was applied to the entire data set. The Cellpose model missed reproducibly round, very bright nuclei (blue arrow), and StarDist predicted many false-positive cells (right panel).

In the DSB, most algorithms performed better on images classified as small or large fluorescent, compared to images classified as ‘purple tissue’ or ‘pink and purple’ tissue. We used methylene blue-stained skeletal muscle sections as a sample dataset for tissue stained with a single dye and Hematoxylin and eosin (HE) stained kidney paraffin sections as an example for multi-dye stained tissue. Analysis of tissue sections might be compromised by heterogenous image quality cause e.g. by artifacts created at the junctions of tiles. To account for these artifacts all workflows used the fused image as input to the analysis pipeline.

While most nuclei in the skeletal muscle dataset are well separated, some form dense clusters, others are out of focus (**Figure 7A**). The size of nuclei varies ten-fold; their shape ranges from elongated to round. The same preprocessing and model as described for the epithelial cells dataset were used; the Cellpose scale-factor range [0.2, 0.4, 0.6] was explored. [Median 3×3 & invert image] preprocessing combined with the Cellpose nuclei 0.6 model produced satisfactory results without further optimization (**Figure 7B**). Outlines were well defined, some objects were missed, few over-segmented. Neither StarDist nor the U-Net performed similarly well. We could not overcome this limitation by adaptation of preprocessing or binning. Processing of the entire dataset identified inadequate segmentation of dense clusters (**Figure 7C**, white arrow) and occasional over-segmentation of large, elongated nuclei (**Figure 7C**, orange arrow) as the main limitations. Nuclei that are out-of-focus were frequently missed (**Figure 7C**, blue arrow). Limiting the analysis to in-focus nuclei is feasible.

**Figure 7.**
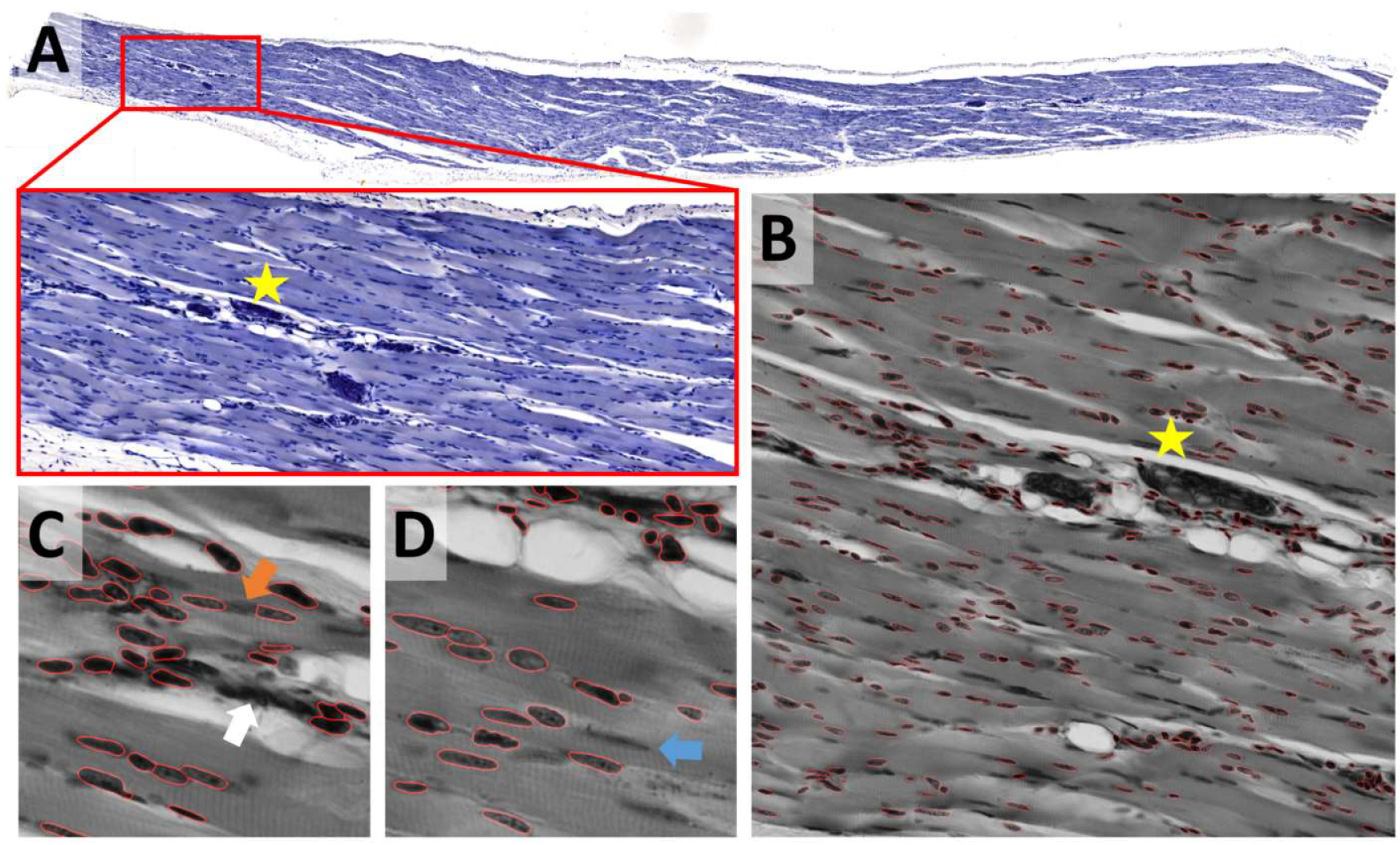
Skeletal muscle notebook: Segmenting irregular nuclei in tissue **(A)** A methylene blue-stained skeletal muscle section was used to test the segmentation of tissue that has been stained with one dye. **(B)** Segmentation was tested on 2048×2048 pixels large image patches. The star shown in **A** and **B** is located at the same position within the image displayed at different zoom factors. The segmentation result (red line) of the Cellpose nuclei 0.6 model is super-imposed onto the original image subjected to [median 3×3] preprocessing. **(C,D)** Close-up on regions that were difficult to segment. Segmentation of dense clusters (white arrow) often failed, and occasional over-segmentation of large, elongated nuclei (orange arrow) was observed. Nuclei that are out-of-focus (blue arrow) were frequently missed (blue arrow).

Cell density is very heterogeneous in the kidney dataset. The Eosin signal from a HE stained kidney paraffin section (**Figure 8A,B**) was obtained by color deconvolution. Nuclei are densely packed within glomeruli and rather sparse in the proximal and distal tubules. Two stitched images were split using OpSeF’s ‘to tiles’ function. Initial optimization was performed on a batch of 16 image tiles, the entire dataset contains 864 tiles. The same preprocessing and model were used as described for the skeletal muscle dataset, the Cellpose scale-factor range [0.6, 0.8, 1.0, 1.4, 1.8] was explored. [Median 3×3 & histogram equalization] preprocessing in combination with the Cellpose nuclei 0.6 model produced fine results (**Figure 8C**). [Mean 7×7 & histogram equalization] preprocessing in combination with StarDist performed similarly well (**Figure 8C**). The latter pipeline resulted in more false-positive detections (**Figure 8C**, purple arrows). The U-Net performed worse, and more nuclei were missed (**Figure 8C**, blue arrow). All models failed for dense cell clusters (**Figure 8C,D**, white arrow).

**Figure 8.**
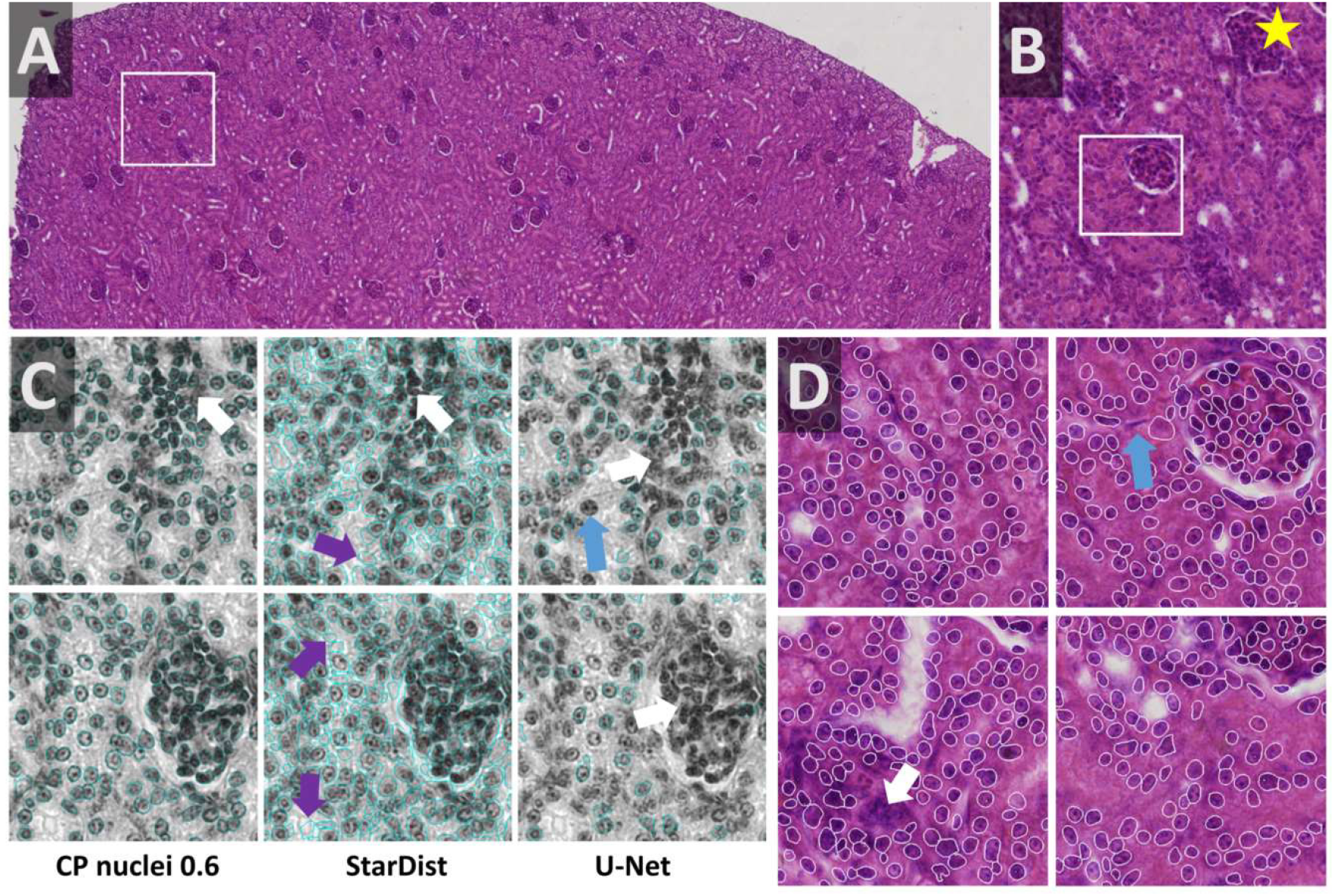
Kidney notebook: Segmenting cell cluster in tissues **(A,B)** Part of a HE stained kidney paraffin section used to test segmentation of tissue stained with two dyes. The white box in **A** highlights the region shown enlarged in **B**. Star in **A,B,D** marks the same glomerulus. **(C)** Eosin signal was extracted by color deconvolution. The segmentation result (blue line) is superimposed onto the original image subjected to [median 3×3] preprocessing. Cellpose nuclei 0.6 model with scale-factor 0.6 in combination with [median 3×3 & histogram equalization] preprocessing and the StarDist model with [mean 7×7 & histogram equalization] performed similarly well **(C,D)**. StarDist resulted in more false-positive detections (purple arrows). The U-Net performed worse, more nuclei were missed (blue arrow). All models failed in very dense areas (white arrow).

Next, we sought to expand the capability of OpSeF to volume segmentation. To this aim, we pre-trained a StarDist 3D model using the annotation of *Arabidopsis thaliana* lateral root nuclei dataset provided by Wolney and colleagues (Wolny et al., 2020). Images were obtained on a Luxendo MuVi SPIM light-sheet microscope (Wolny et al., 2020). We first tested the model with a similar, publically available dataset. Valuchova and colleagues studied differentiation within *Arabidopsis* flowers (Valuchova et al., 2020). To this aim, the authors obtained eight views of H2B:mRuby2 labeled somatic nuclei on a Zeiss Z1 light-sheet microscope. We used a single view to mimic a challenging 3D segmentation (**Figure 9A-C**). Changes in image quality along the optical axis, in particular, deteriorating image quality deeper in the tissue (**Figure 9C**) are a major challenge for any segmentation algorithm. While the segmentation quality of the interactive H-Watershed ImageJ plugin (Vincent and Soille, 1991; Najman and Schmitt, 1996; Lotufo and Falcao, 2000; Schindelin et al., 2015), a state of the art traditional image processing method, is still acceptable in planes with good contrast (**Figure 9B, xy Slice**), results deeper in the tissue are inferior to CNN-based segmentation (data not shown). The H-Watershed plugin consequently fails to segment precisely in 3D (**Figure 9C, zy-slices**). The StarDist 3D model, which was trained on a similar dataset, performs slightly better than the Cellpose nuclei model. To evaluate the performance of these models further, we used the DISCEPTS dataset (Blin et al., 2019). DICEPTS stands for ‘DifferentiatingStemCells & Embryos are a PainToSegment’ and contains various datasets of densely packed nuclei that are heterogeneous in shape, size, or texture. Blin and colleagues elegantly solved the segmentation challenge by labeling the nuclear envelope. We thought to assess the performance of models contained in OpSeF on the more challenging DAPI signal. While the StarDist model trained on *Arabidopsis thaliana* lateral root nuclei shows satisfactory performance on E3.5 mouse blastocysts, notably, the size of nuclei is underestimated, and cells in dense clusters are sometimes missed. Fine-tuning of the non-maximum suppression and the detection threshold might suffice to obtain more precise segmentation results. Interestingly, the more versatile Cellpose cyto, rather than the Cellpose nuclei model, is ideally suited for segmenting the E3.5 mouse blastocysts nuclei (**Figure 9D,E**). Next, we used the neural ‘monolayer’ dataset (**Figure 9D,E**). In this dataset flat cells form tight 3D clusters. It proved to be challenging to segment (Blin et al., 2019). Our pre-trained StarDist model failed to give satisfactory segmentation results (data not shown). The presented cells contain – in contrast to *Arabidopsis thaliana* lateral root nuclei – strong texture in their nuclei. The more versatile Cellpose nuclei model displayed promising initial results with little fine-tuning (**Figure 9D,E**) that might be further improved by re-training the model on the appropriate ground truth.

**Figure 9.**
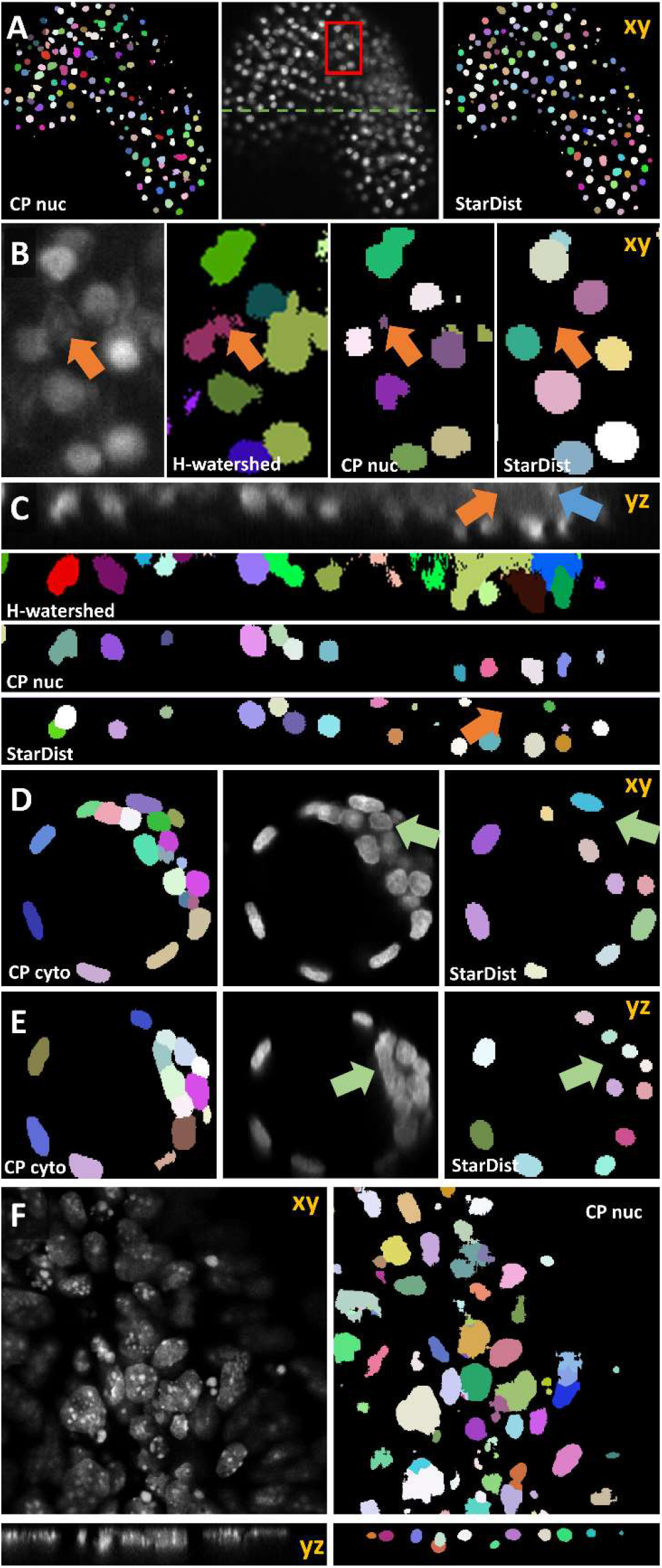
Volume segmentation. In all images, segmented nuclei are assigned a unique random color. **(A)** Side-by-side comparison of segmented somatic nuclei of *Arabidopsis thaliana* flowers obtained with the Cellpose 3D nuclei model (left) and a StarDist 3D model that was trained on a similar dataset (right). The middle panel shows the central xy-plane of the image volume obtained by light-sheet microscopy. The red box marks the area that is shown enlarged in **(B)**. The green line indicates the location of the yz-plane shown in **(C)**. In **(B,C)** the results obtained manually using the interactive ImageJ H-watershed plugin are additionally shown. Orange arrows point at non-nuclear artifact signal that is ignored by StarDist 3D model trained on a dataset containing similar artifacts. Blue arrows point at nuclear signal in a low-contrast area. **(D-F)** Evaluation of 3D model based on the DISCEPTS dataset. Central xy- or yz-planes of original images (grey) are shown side-by-side to the segmentation results of the model as indicated. All data were obtained on confocal microscopes. **(D,E)** Segmentation of E3.5 mouse blastocysts. The StarDist 3D model trained on well-spaced nuclei missed nuclei in dense-cluster (green arrow) that were reliably identified by the Cellpose 3D cyto model. **(H)** Segmentation results of Cellpose nuclei model (right panel) on the neural ‘monolayer’ dataset (left). These cells contain – in contrast to *Arabidopsis thaliana* lateral root nuclei – strong texture in their nuclei, are densely spaced and vary in size.

## Discussion

### Intended use and future developments

Examining the relationship between biochemical changes and morphological alterations in diseased tissues is crucial to understand and treat complex diseases. Traditionally, microscopic images are inspected visually. This approach limits the possibilities for the characterization of phenotypes to more obvious changes that occur later in disease progression. The manual investigation of subtle alterations at the single-cell level, which often requires quantitative assays, is hampered by the data volume. A whole slide tissue image might contain over one million cells. Despite the improvement in machine learning technology, completely unsupervised analysis pipelines have not been widely accepted. Thus, one of the major challenges for the coming years will be the development of efficient strategies to keep the human-expert in the loop. Many biomedical users still perceive deep learning models as black boxes. The mathematical foundation of how CNNs make decisions is improving. OpSeF facilitates understanding the strength of pre-trained models and network architecture on the descriptive, operational level. Thereby, awareness of intrinsic limitations such as the inability of StarDist to segment convex shapes well, or issues relating to the limited field-of-view of neural networks can be reached. It further allows us to quickly assess how robust models are against artifacts such as shadows present in light-sheet microscopy or how well they are in predicting cell shapes accurately, that are neither round nor ellipsoid shaped (e.g. neurons, amoebas). Collectively, increased awareness of limitations and better interpretability of results will be pivotal to increase the acceptance of machine learning methods. It will improve the quality control of results and allow efficient integration of expert knowledge in analysis pipelines (Holzinger et al., 2019a; Holzinger et al., 2019b).

As illustrated in the provided Jupyter notebooks, the U-Net often performed worst. Why is that the case? As previously reported the learning capacity of a single, regular U-Net is limited (Caicedo et al., 2019). Thus, the provision of a set of U-Nets trained on diverse data might be a promising approach to address this limitation (Caicedo et al., 2019), especially in combination with dedicated pre- and post-procession pipelines. OpSeF allows for the straightforward integration of a large number of pre-trained CNNs. We hope that OpSeF will be widely accepted as a framework through which novel models might be made available to other image analysts in an efficient way.

OpSeF allows semi-automated exploration of a large number of possible combinations of preprocessing pipelines and segmentation models. Even if sufficient results are not achievable with pre-trained models, OpSeF results may be used as a guide for which CNN architecture, re-training on manually created labels might be promising. The generation of training data is greatly facilitated by a seamless integration in ImageJ using the AnnotatorJ plugin. We hope that many OpSeF users will contribute their training data to open repositories and will make new models available for integration in OpSeF. Thus, OpSeF might soon become, an interactive model repository, in which an appropriate model might be identified with reasonable effort. Community provided Jupyter notebooks might be used to teach students in courses how to optimize CNN based analysis pipelines. This could educate them and make them less dependent on turn-key solutions that often trade performance for simplicity and offer little insight into the reasons why the CNN-based segmentation works or fails. The better users understand the model they use, the more they will trust them and, the better they will be able to quality control them.

### Integrating various segmentation strategies and quality control of results

Multiple strategies for instance segmentation have been pursued. The U-Net belongs to the ‘pixel to object’ class of methods: each pixel is first assigned to a semantic class (e.g., cell or background), then pixels are grouped into objects (Ronneberger et al., 2015). Mask R-CNNs belong to the ‘object to pixel’ class of methods (He et al., 2017): the initial prediction of bounding boxes for each object is followed by a semantic segmentation. Following an intermediate approach, Schmidt and colleagues first predict star-convex polygons that approximate the shape of cells and use non-maximum suppression to prune redundant predictions (Schmidt et al., 2018b; Weigert et al., 2019). Stringer and colleagues use stimulated diffusion originating from the center of a cell to convert segmentation masks into flow fields. The neural network is then trained to predict flow fields, which can be converted back into segmentation masks (Stringer et al., 2020). Each of these methods has specific strengths and weaknesses. The use of flow fields as auxiliary representation proved to be a great advantage for predicting cell shapes that are not roundish. At the same time, Cellpose is the most computationally demanding model used. In our hands, Cellpose tended to result in more obviously erroneously missed objects, in particular, if objects displayed a distinct appearance compared to their neighbors (blue arrows in **Figure 5B, Figure 6B**, and **Figure 7D**). StarDist is much less computationally demanding, and star-convex polygons are well suited to approximate elliptical cell shapes. The pre-trained StarDist model implemented in OpSeF might be less precise in predicting other convex shapes it has not been trained on. However, this limitation might be overcome by re-training. Many shapes e.g. maple leaves (**Figure 4C**), are concave, and StarDist – due to ‘limitation by design’ – cannot be expected to segment these objects precisely. Segmentation errors by the StarDist model were generally plausible. It tended to predict cell-like shapes, even if they are not present (**Figure 6B**). Although the tendency of StarDist to fail gracefully might be advantageous in most instances, this feature requires particularly careful quality control to detect and fix errors. The ‘pixel-to-object’ class of methods is less suited for segmentation of dense cell clusters. The misclassification of just a few pixels might lead to the fusion of neighboring cells.

OpSeF integrates three mechanistically distinct methods for CNN-based segmentation in a single framework. This allows comparing these methods easily. Nonetheless, we decided against integrating an automated evaluation, e.g. by determining F1 score, false positive and false negative rates, and accuracy. Firstly, for most projects no ground-truth is available. Secondly, we want to encourage the user to visually inspect segmentation results. Inspecting 100 different segmentation results opened in ImageJ as stack takes only a few minutes and gives valuable insight into when and how segmentations fail. This knowledge is easily missed when just looking at the output scores of commonly used metrics but might have a major impact on the biological conclusion. Even segmentation results from a model with 95% precision and 95% recall for the overall cell population might be not suited to determine the abundance of a rare cell type if these cells are systematically missed, detected less accurately in the mutant situation, or preferentially localized to areas in the tissue that are not segmented well. Although it is difficult to capture such issues with common metrics, they are readily observed by a human expert. Learning more about the circumstances in which certain types of CNN-based segmentation fail helps to decide when human experts are most essential for quality control of results. Moreover, it is pivotal for the design of post-processing pipelines. These might select among multiple segmentation hypotheses – on an object by object basis – the one, which gives the most consistent results for reconstructing complex cells-shapes in large 3D volumes or for cell-tracking.

### Optimizing results and computational cost

Image analysis pipelines are generally a compromise between ease-of-use and performance as well as between computational cost and accuracy. Until now, rather simple, standard U-Nets are – despite their limited learning capacity – most frequently used models in the major image analysis tools. In contrast, the winning model of the 2018 Data Science Bowl by the [ods.ai] topcoders team used sophisticated data post-processing to combine the output of 32 different neural networks (Caicedo et al., 2019). The high computational cost currently limits the widespread use of this or similar approaches. OpSeF is an ideal platform to find the computationally most efficient solution to a segmentation task. The [ods.ai] topcoders algorithm was designed to segment five different classes of nuclei: ‘small’ and ‘large fluorescent’, ‘grayscale’, ‘purple tissue’ and ‘pink and purple tissue’(Caicedo et al., 2019). Stringer and colleagues used an even broader collection of images that included cells of unusual appearance and natural images of regular cell-like shapes such as shells, garlic, pearls, and stones (Stringer et al., 2020).

The availability of such versatile models is precious, in particular, for users, who are unable to train custom models or lack resources to search for the most efficient pre-trained model. For most biological applications, however, no one-fits-all solution is required. Instead, potentially appropriate models might be pre-selected, optimized, and tested using OpSeF. Ideally, an image analyst and a biomedical researcher will jointly fine-tune the analysis pipeline and quality control results. This way, resulting analysis workflows will have the best chances of being both robust and accurate, and an ideal balance between manual effort, computational cost, and accuracy might be reached.

Comparison of the models available within OpSeF revealed that the same task of segmenting 100 images using StarDist took 1.5-fold, Cellpose with fixed scale-factor 3.5-fold, and Cellpose with flexible scale-factor 5-fold longer compared to segmentation with the U-Net.

The systematic search of the optimal parameter and ideal performance might be dispensable if only a few images are to be processed that can be easily manually curated, but highly valuable if massive datasets produced by slide-scanner, light-sheet microscopes or volume EM techniques are to be processed.

### Deployment strategies

We decided against providing OpSeF as an interactive cloud solution. A local solution uses existing resources best, avoids limitations generated by the down- and upload of large datasets, and addresses concerns regarding the security of clinical datasets. Although the provision of plugins is perceived as crucial to speed up the adoption of new methods, image analysts increasingly use the Jupyter notebooks that allow them to document workflows step-by-step. This is a major advantage compared to interactive solutions, in which parameters used for analysis are not automatically logged. Biologists might hesitate to use Jupyter notebooks for analysis due to an initial steep learning curve. Once technical challenges such as the establishment of the conda environment are overcome, notebooks allow them to integrate data analysis and documentation with ease. Notebooks might be deposited in repositories along with the raw data. This builds more trust in published results by improving transparency and reproducibility.

## Author Contributions

T.M.R. designed and developed the OpSeF. R.H. and P.H. designed and developed the AnnotatorJ integration. Data was analyzed by T.M.R. Manuscript was written by T.M.R. with contributions from all authors.

## Acknowledgments

We would like to thank Thorsten Falk and Volker Hilsenstein (Monash Micro Imaging) for testing of the software and critical comments on the manuscript; the IT service group of the MPI Heart and Lung Research for technical support. The authors gratefully acknowledge Varun Kapoor (Institut Curie), Constantin Pape (EMBL), Carsen Stringer (Janelia Research Campus), and Adrian Wolny (EMBL) for advice. We thank anyone who made their data available, in particular: the Broad Bioimage Benchmark Collection for image set BBBC027 (Svoboda et al., 2011) and BBBC038v1 (Caicedo et al., 2019) and Christoph Möhl.

